# Digital color-coded molecular barcoding reveals dysregulation of common FUS and FMRP targets in soma and neurites of ALS mutant motoneurons

**DOI:** 10.1101/2022.08.02.502510

**Authors:** Maria Giovanna Garone, Debora Salerno, Alessandro Rosa

## Abstract

Mutations in RNA binding proteins (RBPs) have been linked to the motor neuron disease amyotrophic lateral sclerosis (ALS). Extensive auto-regulation, cross-regulation, cooperation and competition mechanisms among RBPs are in place to ensure proper expression levels of common targets, often including other RBPs and their own transcripts. Moreover, several RBPs play a crucial role in the nervous system by localizing target RNAs in specific neuronal compartments. These include the RBPs FUS, FMRP and HuD. ALS mutations in a given RBP are predicted to produce a broad impact on such delicate equilibrium. Here we studied the effects of the severe FUS-P525L mutation on common FUS and FMRP targets. Expression profiling by digital color-coded molecular barcoding in cell bodies and neurites of human iPSC-derived motor neurons revealed altered levels of transcripts involved in cytoskeleton, neural projection and synapses. One of the common targets is HuD, which is upregulated because of the loss of FMRP binding to its 3’UTR due to mutant FUS competition. Notably, many genes are commonly altered upon FUS mutation or HuD overexpression, suggesting that a substantial part of the effects of mutant FUS on the motor neuron transcriptome could be due to HuD gain-of-function. Among altered transcripts we also identified other common FUS and FMRP targets, namely MAP1B, PTEN, and AP2B1, that are upregulated upon loss of FMRP binding on their 3’UTR in FUS-P525L motor neurons. This work demonstrates that the impairment of FMRP function by mutant FUS might alter the expression of several genes, including new possible biomarkers and therapeutic targets for ALS.

## INTRODUCTION

RNA-binding proteins (RBPs) play a key role in controlling RNA metabolism and proteome complexity and functions in neurons, as well as in the fast and localized modulation of neuronal gene expression. These functions are crucial for proper axon and dendrite development, guidance, targeting, and synapse formation. Changes in RBP subcellular localization have been observed in neurodegenerative diseases such as amyotrophic lateral sclerosis (ALS), a disorder primarily caused by the loss of upper and lower motor neurons (MNs) that leads to progressive impairment of the neuromuscular system, axonal degeneration, and neuronal death (Brown and Al-Chalabi, 2017). The alteration of RNA metabolism in MNs is considered one of the overt causes of the onset of ALS, as several RBPs, including FUS, are mutated or aggregated in patients (Lagier-Tourenne et al., 2010). Most ALS-associated FUS mutations are missense mutations mainly localized in the C-terminal nuclear localization signal (NLS), leading to reduced affinity with the nuclear import receptor Transportin-1 (TNPO1) and resulting in protein mislocalization to the cytoplasm (Dormann et al., 2010; Deng et al., 2014; Tyzack et al., 2019).

Although FUS is a ubiquitous protein, it plays specific functions in neuronal cells. Indeed, it localizes in specific compartments such as the neuromuscular junction (NMJ) (So et al., 2018), presynaptic terminals (Schoen et al., 2015; Sahadevan et al., 2022), and post-synaptic dendrites (Aoki et al., 2012; Belly et al., 2005; Fujii et al., 2005; Yasuda et al., 2013), where it acts as local translation regulator (López-Erauskin et al., 2018; Yasuda et al., 2013) and is responsible for synaptic structure and function (Fujii et al., 2005; Fujii and Takumi, 2005).

We previously showed strong correlation between changes in protein levels and selective binding of the P525L mutant FUS, which is highly delocalized in the cytoplasm and associated with juvenile and severe ALS (Dormann et al., 2010), to 3’UTRs in human pluripotent stem cell (iPSC)-derived MNs (De Santis et al., 2019). Genes whose transcripts are bound in the 3’UTR by mutant FUS show altered protein levels and are involved in cytoskeleton and neuron projection (Garone et al., 2020). This suggests that changed targeting by cytoplasmic mutant FUS might promote axonopathy. Indeed, axonal dysfunction has been reported in early symptomatic ALS patients and it occurs before the motor phenotype in animal models (Fischer et al., 2004; Roy et al., 2005), indicating that ALS can be classified as a distal axonopathy (Moloney et al., 2014). ALS axo-pathomechanisms include alteration in the neuronal cytoskeleton, RNA transport, axonal energy supply, clearance of junk proteins, and aberrant axonal branching (Suzuki et al., 2020). To this regard, we recently reported increased axon branching and arborization, as well as faster outgrowth after injury in FUS-ALS MNs, due to increased activity of the neuronal RBP HuD (Garone et al., 2021).

HuD (encoded by the *ELAVL4* gene) is required for neuron development, including differentiation and axonal outgrowth (Robinow et al., 1988; Akamatsu et al., 2005), and only recently has gained attention in the field of neurodegeneration as a component of cytoplasmic inclusions in FUS, TDP-43, and sporadic ALS patients (Blokhuis et al., 2016; De Santis et al., 2019). In addition to contributing to synaptic plasticity in adult neurons, HuD has been correlated to the response of peripheral neurons to damage (Sanna et al., 2016). We have shown that HuD levels are increased in FUS^P525L^ human iPSC-derived MNs and in the spinal cord of a FUS-ALS mouse model (De Santis et al., 2017; Garone et al., 2021). This is determined by the binding of mutant FUS to the 3’UTR of HuD mRNA, which is also a binding target of the translational repressor FMRP (fragile-X mental retardation protein) (De Santis et al., 2019; Garone et al., 2021). In ALS MNs, a pathological mechanism results from the competition between mutant FUS and FMRP for HuD 3’UTR binding. Mutant FUS binding hinders HuD translational repression by FMRP, leading to increased HuD levels with consequences on HuD targets, NRN1 and GAP43, in turn involved in abnormal axonal morphology and recovery upon injury (Garone et al., 2021). Notably, similar axonal phenotypes have been reported in other non-FUS ALS models (Osking et al., 2019; Kabashi et al., 2010), and upregulated levels of HuD have been observed in sporadic ALS patients’ motor cortex (Dell’Orco et al., 2021). These evidences suggest a possible transversal role of HuD in ALS (both sporadic and familial), and its involvement in early axonopathy phenotypes.

In this work, we aimed to gain insight into the molecular mechanisms that could underlie axonal dysfunctions in ALS, with a specific focus on the interplay between three RBPs: ALS mutant FUS, FMRP and HuD. By leveraging human iPSC-derived MNs cultured in compartmented chambers and RNA profiling by digital color-coded molecular barcoding in soma and neurites, we found altered expression of genes involved in neurodevelopment, cytoskeleton and synapse formation upon FUS mutation, FMRP loss or HuD overexpression. Among altered genes, we identified *MAP1B, AP2B1* and *PTEN*, which represent new common FMRP and FUS^P525L^ targets. These genes have been previously involved in several aspects of nerve growth, arborization and regeneration, in physiological and pathological conditions (Park et al., 2008; Koscielny et al., 2018; Villarroel-Campos and Gonzalez-Billault, 2014). Translation of *MAP1B, AP2B1* and *PTEN* is negatively controlled by FMRP in FUS wild-type MNs, while we observed increased levels in FUS mutant MNs. This work suggests that the intrusion of mutant FUS into FMRP functions might exert a broad impact on the transcriptome of both neurites and soma in ALS MNs.

## RESULTS

### Identification of common FMRP and FUS targets

We have previously identified the transcripts bound by FUS in iPSC-derived human MNs, showing that while FUS^WT^ preferentially binds introns, the ALS mutant FUS^P525L^ interacts with 3’UTRs (De Santis et al., 2019). Thus, the consequences of FUS pathological mutations might arise by both loss-of-function (loss of intron binding due to reduced nuclear FUS levels) and gain-of-function (gain of 3’UTR binding due to increased accumulation of FUS in the cytoplasm) mechanisms. Changes in FUS mRNA targets correlated with changes in the proteome (Garone et al., 2020). Moreover, we discovered competition between mutant FUS and FMRP for the binding to HuD 3’UTR, resulting in the loss of negative regulation by FMRP on HuD translation (Garone et al., 2021). These observations prompted us to assess if ALS FUS mutation might interfere with FMRP activities on other relevant genes, including genes that might be affected at the level of alternative splicing by loss of FUS^WT^ and genes targeted in their 3’UTR by FUS^P525L^. The possible common targets of FMRP and FUS were identified from previously published FUS PAR-CLIP (De Santis et al., 2019) and FMRP HITS-CLIP (Darnell et al., 2011) data. We found a common set of 1135 targets bound both by FUS^WT^ in intronic regions and by FUS^P525L^ in the 3’UTR (**Figure 1A**). Cross-reference with FMRP HITS-CLIP data, comprising FMRP interacting mRNAs whose translation is inhibited by delaying ribosomal translocation at a p-value < 0.05, identified 136 common targets (**Figure 1A**) (**Supplementary Table S1**). Gene ontology (GO) term enrichment analysis of this subset revealed categories mainly related to cytoskeletal protein binding (molecular function, GO: MF), neuron projection development and morphogenesis (biological process, GO: BP), neuron projection, plasma membrane bounded cell projection, synapse (cellular component, GO: CC) among the top five terms (**Figure 1B**; **Supplementary Table S2**), suggesting that common FMRP and FUS targets include genes that play crucial roles in subcellular neuronal compartments.

**Figure 1.**
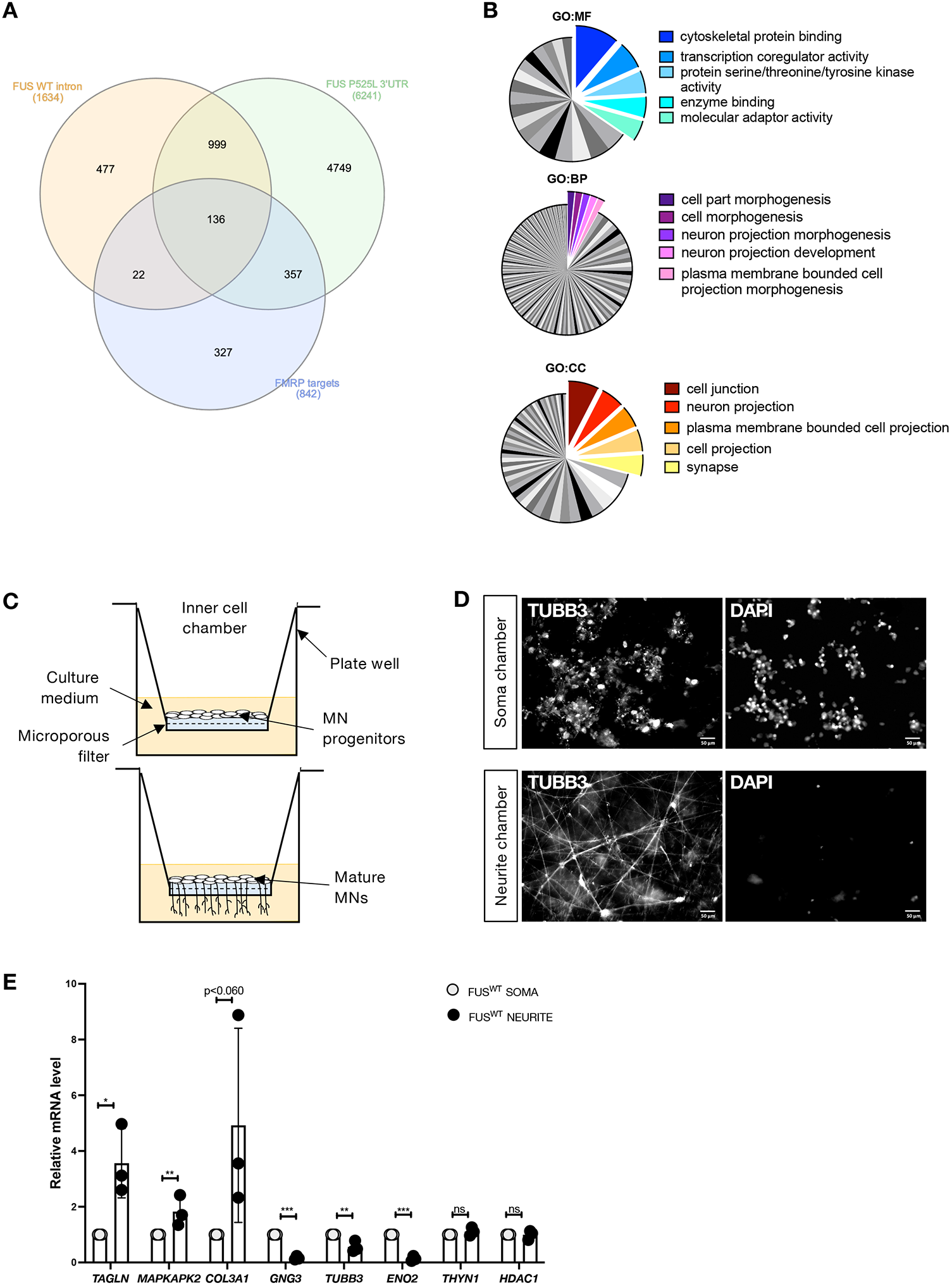
FMRP and FUS interactomics analyses and neuron cell culture system. (**A**) Venn diagram showing the overlap among the transcripts bound in intronic regions by FUS^WT^ (p-value <0.05) and in the 3′UTR by endogenous FUS^P525L^ (p-value <0.05) and FMRP targets (p-value <0.05). (**B**) GO term enrichment analysis of the 136 overlapped genes. The -log10 (adjusted p-value) associated with each category is represented by the size of the pie slice. GO:MF (molecular function); GO:BP (biological process); GO:CC (cellular component). (**C**) Schematic representation of dissociated and re-plated iPSC-derived spinal MN progenitor cells at day 5 of differentiation (top) and mature MNs (bottom) into cell culture insert. (**D**) Immunostaining of TUBB3 and staining with DAPI in MNs cultured in cell culture insert. Scale bar: 50 µm. (**E**) Analysis of the mRNA levels of the indicated genes by real-time qRT-PCR in iPSC-derived spinal MNs. *TAGLN, MAPKAPK2, COL3A1* are neurite markers; *GNG3, ENO2* are neuronal cell body enriched markers; *THYN1, HDAC1* are housekeeping genes, included in this analysis as uniformly distributed transcripts. The graph shows the average from three independent differentiation experiments, error bars indicate the standard deviation (Student’s t-test; paired; two tails; *p < 0.05; **p < 0.01; ***p < 0.001; n.s. non significant).

Taken together these data suggest that the FUS mutation might impinge the expression levels, splicing and/or subcellular localization of several FMRP mRNA targets. This evidence prompted us to analyze the levels of common FMRP and FUS mRNA interactors in soma and neurite compartments.

### Inner chamber iPSC-derived MN cultures to analyze gene expression in soma and neurites

In order to isolate the RNA from different cellular compartments without cross-contamination, we took advantage of modified Boyden chambers, which have been previously used to study the axonal transcriptome of mouse embryo dorsal root ganglia (Minis et al., 2014). Human iPSCs were converted into spinal MNs by ectopic expression of a transcription factor module, including *Ngn2, Isl1*, and *Lhx3* (NIL), with a protocol that allows the production of nearly pure motoneuronal population without the need of cell sorting (De Santis et al., 2018; Garone et al., 2019). MN progenitors at day 5 of differentiation were dissociated and re-plated inside the inner chamber onto a porous membrane that enabled axons to grow across the filter, restricting the cell bodies to the top membrane surface (**Figure 1C**). Correct compartmentalization of soma and neurites in the two cell culture sides was assessed by immunostaining of the neuronal tubulin, TUBB3, and labelled nuclei with DAPI in iPSC-derived MNs at day 12 of differentiation in cell culture inserts (**Figure 1D**). RNA samples collected from soma and neurites were then analyzed by quantitative RT-PCR. We observed enrichment of the well-characterized neuronal projections markers *TAGLN, COL3A1* and *MAPKAK2* in the neurite compartment, and enrichment of *GNG3* and *ENO2* in the soma (**Figure 1E**), consistent with the known localization of these transcripts in neurons (Ludwik et al., 2019).

### Single-mRNA molecules detection of FMRP and FUS targets in soma and neurites

The 136 FUS and FMRP common targets were examined based on their known role in neurodevelopmental and/or neurodegenerative disorders by taking advantage of the DISEASES database (Pletscher-Frankild et al., 2015). 70 candidates with the highest z-score (**Supplementary Table S3**) were selected for gene expression analysis in soma and axonal compartments. In consideration of previous findings (Garone et al., 2021), we added to this list HuD (*ELAVL4*) and its target genes, *NRN1* and *GAP43*. Transcripts quantification was performed by NanoString digital color-coded molecular barcoding. This technology employs fluorescent barcodes that allow for the direct detection of distinct mRNA targets in a single run without amplification, with a sensitivity of 1 copy per cell and requiring nanoscale amounts of RNA (Salerno and Rosa, 2020). A code set specific to a 100-base region of the target mRNA was designed using a 3′ biotinylated capture probe and a 5′ reporter probe tagged with a specific fluorescent barcode, creating two sequence-specific probes for each target transcript (**Supplementary Table S4**).

MNs differentiated from an isogenic pair of FUS^WT^ and FUS^P525L^ iPSCs (Lenzi et al., 2015) were cultured in modified Boyden chambers to allow RNA isolation from soma and neurites. We identified 24 and 22 targets that were differentially expressed in FUS^P525L^ motoneurons at p-value <0.05, respectively in the soma and neurite compartment (**Figure 2A-B**). Interestingly, over 80% of gene expression changes consisted in overexpression. We performed GO enrichment analysis to characterize the biological process and cellular component of these genes (**Figure 2C, Supplementary Table S5**). We found that targets altered in the neurite subcellular compartment showed enrichment in biological processes involving “regulation of trans-synaptic signaling”, “transport along microtubule”, “cytoskeleton-dependent intracellular transport”, and cellular components related to “synapse”, “neuron projection”, “cell junction” and “dendrite” (**Figure 2C**). On the other hand, altered targets in the cell body fraction are involved in nervous “system development”, “neuron projection morphogenesis”, “neuron differentiation”, and enriched for “cell junction”, “postsynapse” and “neuron projection” compartment (**Figure 2C**). Together, these data suggest that the FUS mutation might impact the synaptic component in axons and affect the function of genes in the soma involved in neuron development and morphogenesis. We then performed the same analysis in MNs derived from isogenic FMRP^WT^ and FMRP^KO^ human iPSCs (Brighi et al., 2021). We observed 42 differentially expressed targets in the soma and 18 in neurites in FMRO^KO^ MNs at a p-value < 0.05 (**Figure 3A-B**). For these genes, GO enrichment analysis revealed categories related to the regulation of transport and localization, and regulation of cellular processes and development in both soma and neurite domain (**Figure 3C, Supplementary Table S5**). These data support the role of FMRP as an essential RBP for transporting mRNAs to subcellular neuron sites with implication in neurodevelopmental diseases.

**Figure 2.**
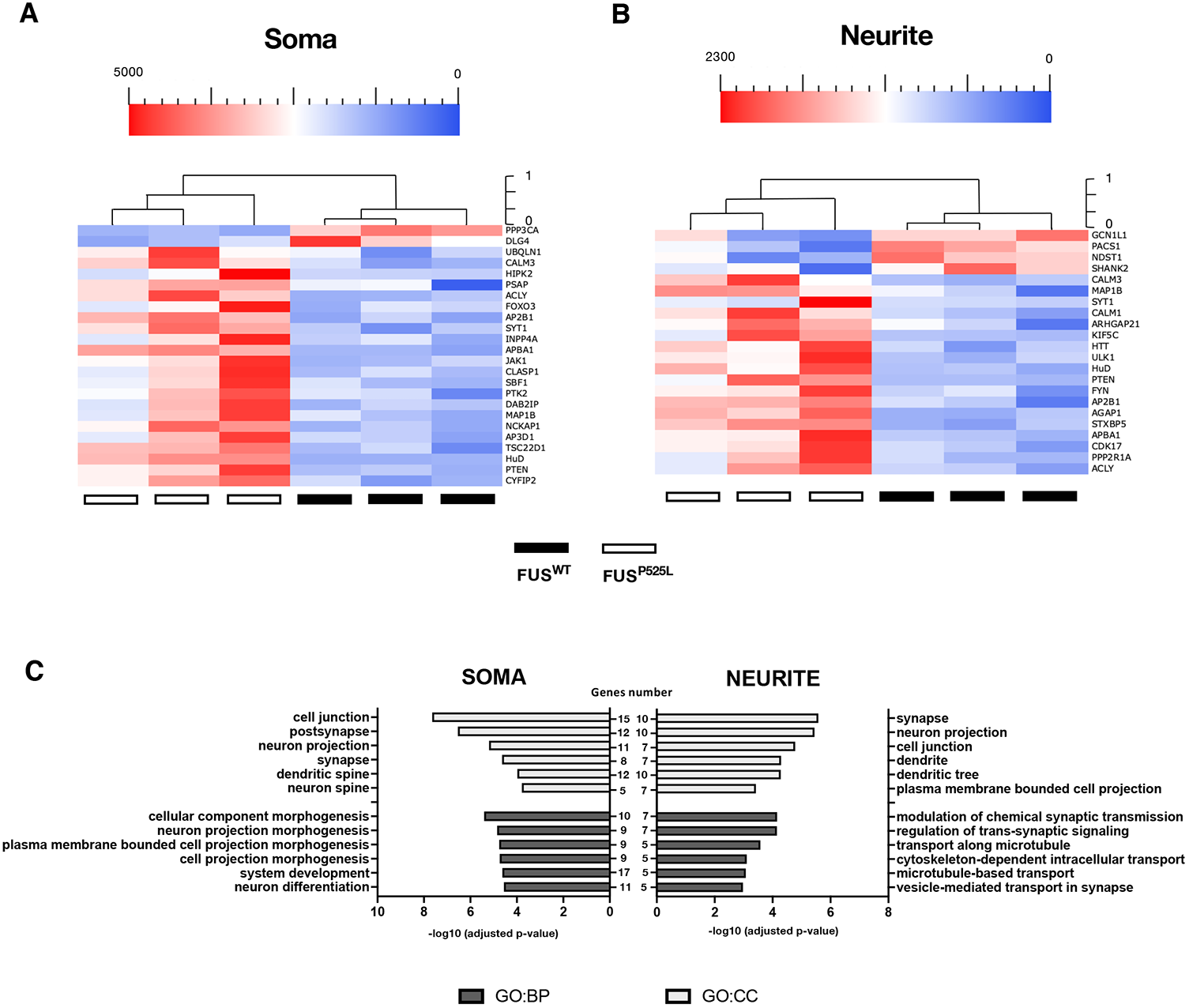
Differentially expressed genes in soma and neurite of FUS^WT^ and FUS^P525L^ MNs. (**A**,**B**) Heatmap representing the RNA molecules count of differentially expressed genes in the cell body (A) or neurite (B) compartment of iPSC-derived FUS^WT^ and FUS^P525L^ MNs. Plotted values correspond to normalized RNA molecules as described in the Methods section. Red, higher abundance; blue, lower abundance. (**C**) Bar graph representation of the top 6 GO terms enriched in differentially expressed genes as in (A,B). The -log10(adjusted p-value) and number of genes are shown.

**Figure 3.**
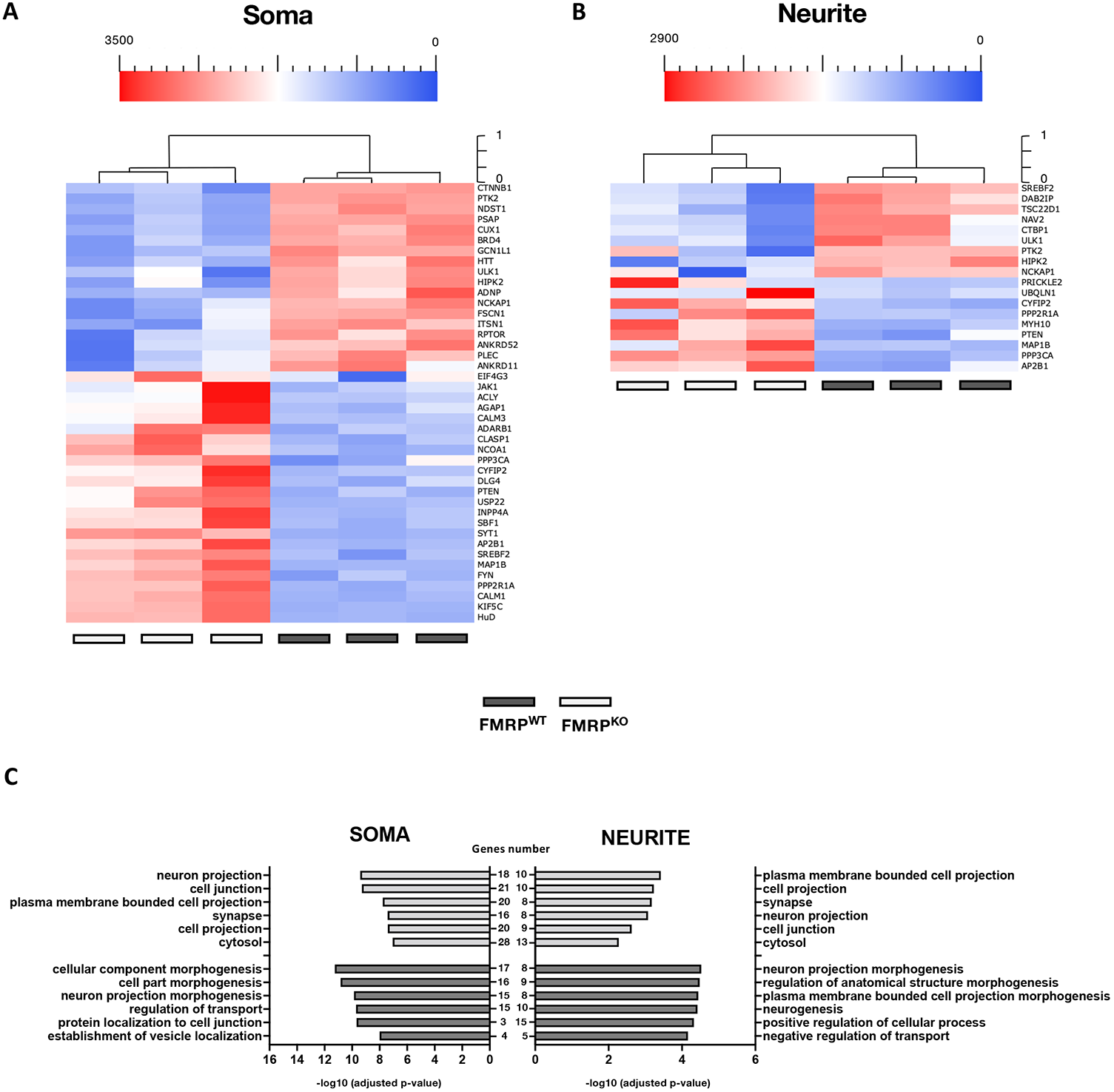
Differentially expressed genes in soma and neurite of FMRP^WT^ and FMRP^KO^ MNs. (**A**,**B**) Heatmap representing the RNA molecules count of differentially expressed genes in the cell body (A) or neurite (B) compartment of iPSC-derived FMRP^WT^ and FMRP^KO^ MNs. Plotted values correspond to normalized RNA molecules as described in the Methods section. Red, higher abundance; blue, lower abundance. (**C**) Bar graph representation of the top 6 GO terms enriched in differentially expressed genes as in (A,B). The -log10(adjusted p-value) and number of genes are shown.

### Increased HuD levels in MNs mimics the effects of ALS mutant FUS

One of the consequences of the impaired FMRP activity in FUS mutant MNs is the upregulation of HuD. Notably, we found increased HuD levels in neurites and soma of FUS^P525L^ and in the soma of FMRP^KO^ MNs (**Figures 2-3**), confirming our previous observation (Garone et al., 2021). Thus, changes in the transcriptome in both these genetic backgrounds might be due, to some extent, to HuD increased levels. To experimentally address this possibility, we performed single-mRNA molecules detection in HuD overexpressing MNs, with an otherwise FUS^WT^ and FMRP^WT^ background. To this aim, we took advantage of a stably transfected iPSC line expressing a HuD transgene under the control of the neuronal-specific human synapsin 1 promoter (SYN1::HuD). Importantly, in iPSC-derived SYN1::HuD MNs, HuD levels were increased in a range similar to that observed in FUS^P525L^ or FMRP^KO^ cells (Garone et al., 2021). 56 genes in the soma and 14 genes in the neurites were differentially expressed upon HuD overexpression (**Figure 4A-B**). Also in this case, most of the change was represented by upregulation. GO enrichment analysis showed an interesting overlap with categories resulting from the analysis of FUS^P525L^ MNs. Indeed, altered neurite transcripts were involved in biological processes related to synaptic transmission. On the other hand, neuronal morphogenesis and differentiation terms were enriched in the soma (**Figure 4C, Supplementary Table S5**). In agreement with this observation, analysis of differentially expressed transcripts showed that SYN1::HuD biological replicates tend to cluster with FUS^P525L^ samples, rather than with their common parental line (FUS^WT^), both in the soma and neurite compartments (**Figure 5**).

**Figure 4.**
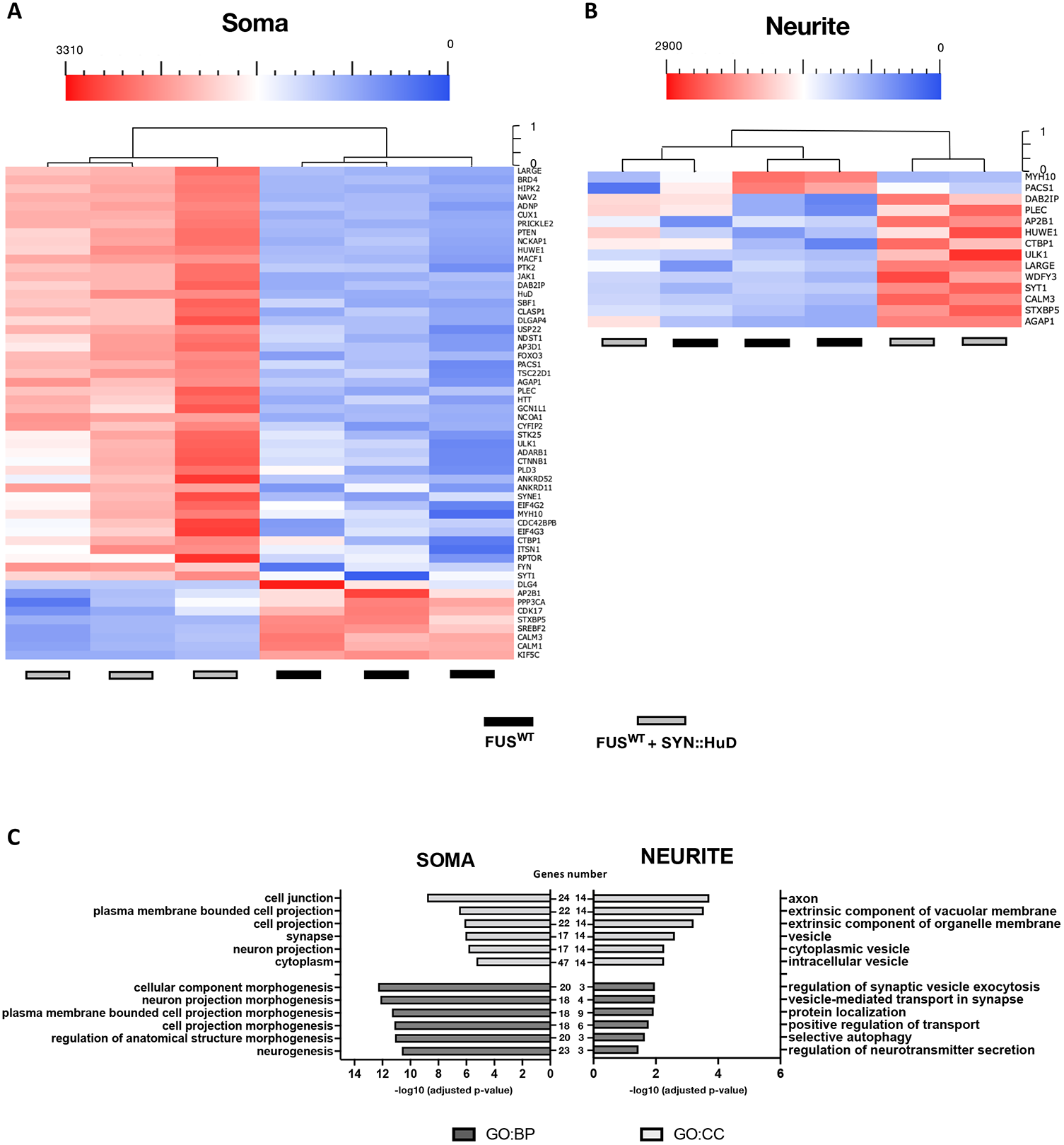
Differentially expressed genes in soma and neurite of FUS^WT^ and FUS^WT^ SYN::HuD MNs. (**A**,**B**) Heatmap representing the RNA molecules count of differentially expressed genes in the cell body (A) or neurite (B) compartment of iPSC-derived FUS^WT^ MNs and MNs derived from a stably transfected FUS^WT^ iPSC line expressing a HuD transgene under the control of the neuronal-specific human synapsin 1 promoter (SYN1::HuD). Plotted values correspond to normalized RNA molecules as described in the Methods section. Red, higher abundance; blue, lower abundance. (**C**) Bar graph representation of the top 6 GO terms enriched in differentially expressed genes as in (A,B). The - log10(adjusted p-value) and number of genes are shown.

**Figure 5.**
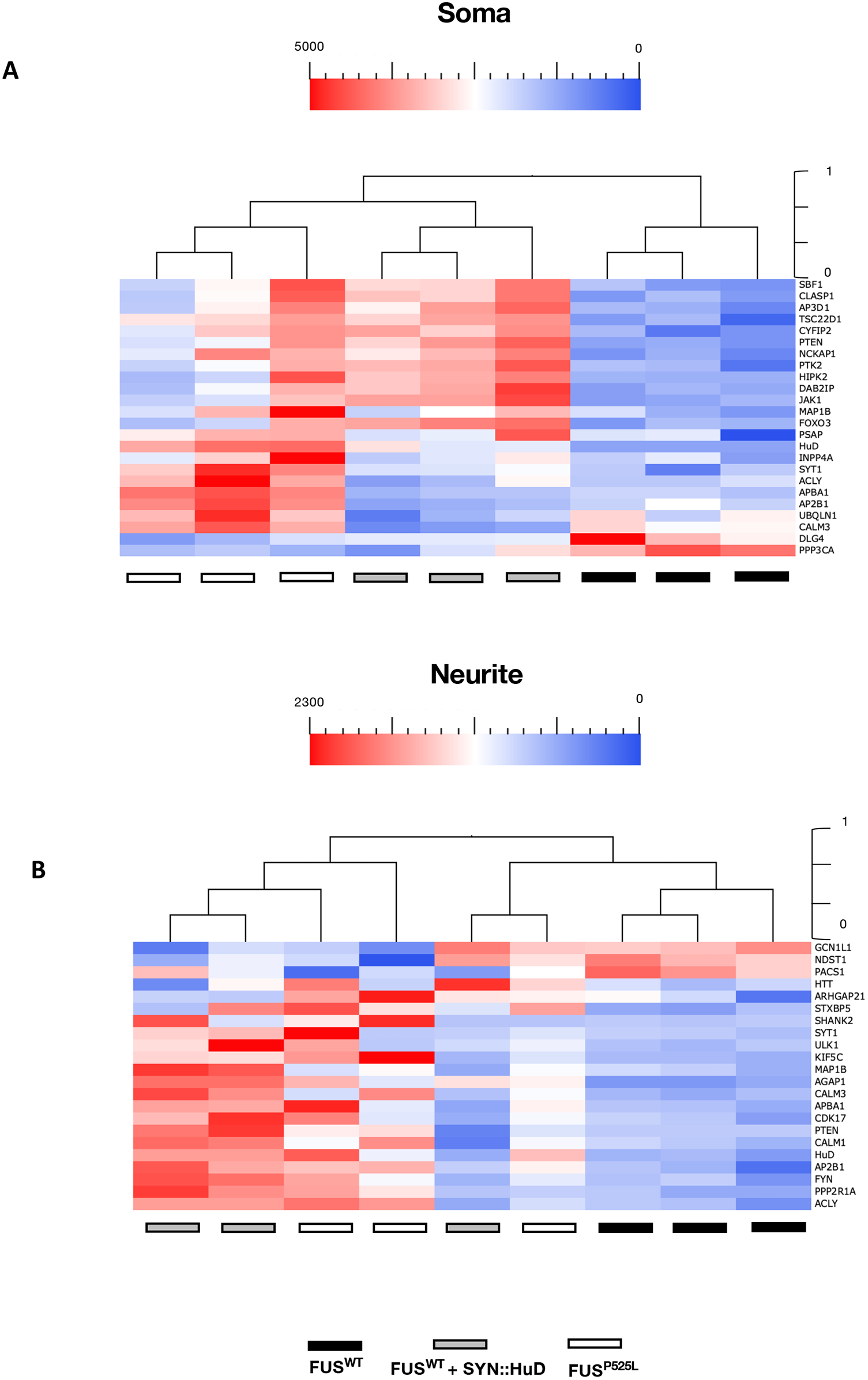
Differentially expressed genes in soma and neurite of FUS^WT^, FUS^P525L^ and FUS^WT^ SYN::HuD MNs. (**A**,**B**) Heatmap representing the RNA molecules count of differentially expressed genes in the cell body (A) or neurite (B) compartment of iPSC-derived FUS^WT^, FUS^P525L^ and FUS^WT^ SYN::HuD MNs. Plotted values correspond to normalized RNA molecules as described in the Methods section. Red, higher abundance; blue, lower abundance.

Collectively, these results suggest that increased HuD levels might substantially contribute to altered gene expression observed in FUS mutant MNs, in particular for disease-relevant transcripts possibly involved in axopathogenesis.

### ALS mutant FUS competes with FMRP for 3’UTR binding of disease relevant genes

We next focused on common FUS and FMRP targets that, similarly to HuD, were increased in both FUS^P525L^ and FMRP^KO^ MNs and identified three interesting candidates. *AP2B1* (adaptor related protein complex 2 subunit beta 1) encodes for a subunit of the adaptor protein complex 2 (AP2), which promotes protein transport via transport vesicles in several membrane traffic pathways. Adaptor protein complexes are parts of the vesicle coat that is involved in cargo selection and vesicle assembly (McMahon and Boucrot, 2011). Moreover, AP2 assists the recycle of synaptic vesicle membranes from the presynaptic surface (De Camilli and Takei, 1996). *MAP1B*, a well-known FMRP target, plays an important role in the tyrosination of alpha-tubulin in neuronal microtubules and is involved in the cytoskeletal modifications associated with neurite outgrowth (Sayas and Ávila, 2014). *PTEN* (phosphatase and tensin homolog) encodes for a protein phosphatase that has been recently implicated in neurodevelopmental and neurodegenerative diseases (Rademacher et al., 2019; Cummings et al., 2022; Kirby et al., 2011; Little et al., 2015; Stopford et al., 2017).

FMRP binding to these transcripts in the presence or absence of mutant FUS was evaluated by native RNA immunoprecipitation (RIP). FMRP was immunoprecipitated from human iPSC-derived MN extracts, and the associated mRNAs were analyzed by quantitative RT-PCR. As previously shown for *HuD* and *MAP1B* (Garone et al., 2021), we observed a significant enrichment of *AP2B1* and *PTEN* in FMRP RIP from FUS^WT^ MNs, while the housekeeping gene ATP5O served as a negative control. Interestingly, FMRP binding to both targets was completely abolished in FUS^P525L^ MNs (**Figure 6A**), similarly to what previously shown for *HuD* and *MAP1B* (Garone et al., 2021). These data suggest that impairment of FMRP binding to target RNAs in mutant FUS MNs, possibly due to competition for 3’UTR binding, might occur for several common FUS and FMRP targets. To assess the functional consequences of the loss of FMRP interaction in the presence of FUS^P525L^, we took advantage of a reporter assay based on a luciferase gene fused to the 3’UTR of the target of interest. *AP2B1, MAP1B* and *PTEN* 3’UTR reporters were transfected in HeLa cells ectopically expressing a wild-type or P525L mutant FUS transgene (both fused to RFP, or RFP alone as control) in combination with a FMRP transgene or eGFP as control. As shown in **Figure 6B**, when FMRP was overexpressed in the presence of RFP alone, the 3′UTR reporter activity was significantly decreased for all the candidates. Notably, co-expression of RFP-FUS^P525L^, but not RFP-FUS^WT^, partially reversed (for *MAP1B*) or completely abolished (for *AP2B1* and *PTEN*) such negative regulation by FMRP. These results suggest that FMRP could act as a negative regulator of *AP2B1, MAP1B* and *PTEN* translation by direct 3’UTR binding, while mutant FUS competition would impair such function. We next analyzed protein levels of all three candidates in iPSC-derived MNs, observing significant up-regulated levels in mutant FUS as well as FMRP KO cells (**Figure 7A, Figure S1**). Importantly, AP2B1, MAP1B and PTEN protein levels were analyzed in MNs derived from an independent isogenic pair of wild-type and P525L iPSC lines (Marrone et al., 2018), confirming this result (**Figure 7B**).

**Figure 6.**
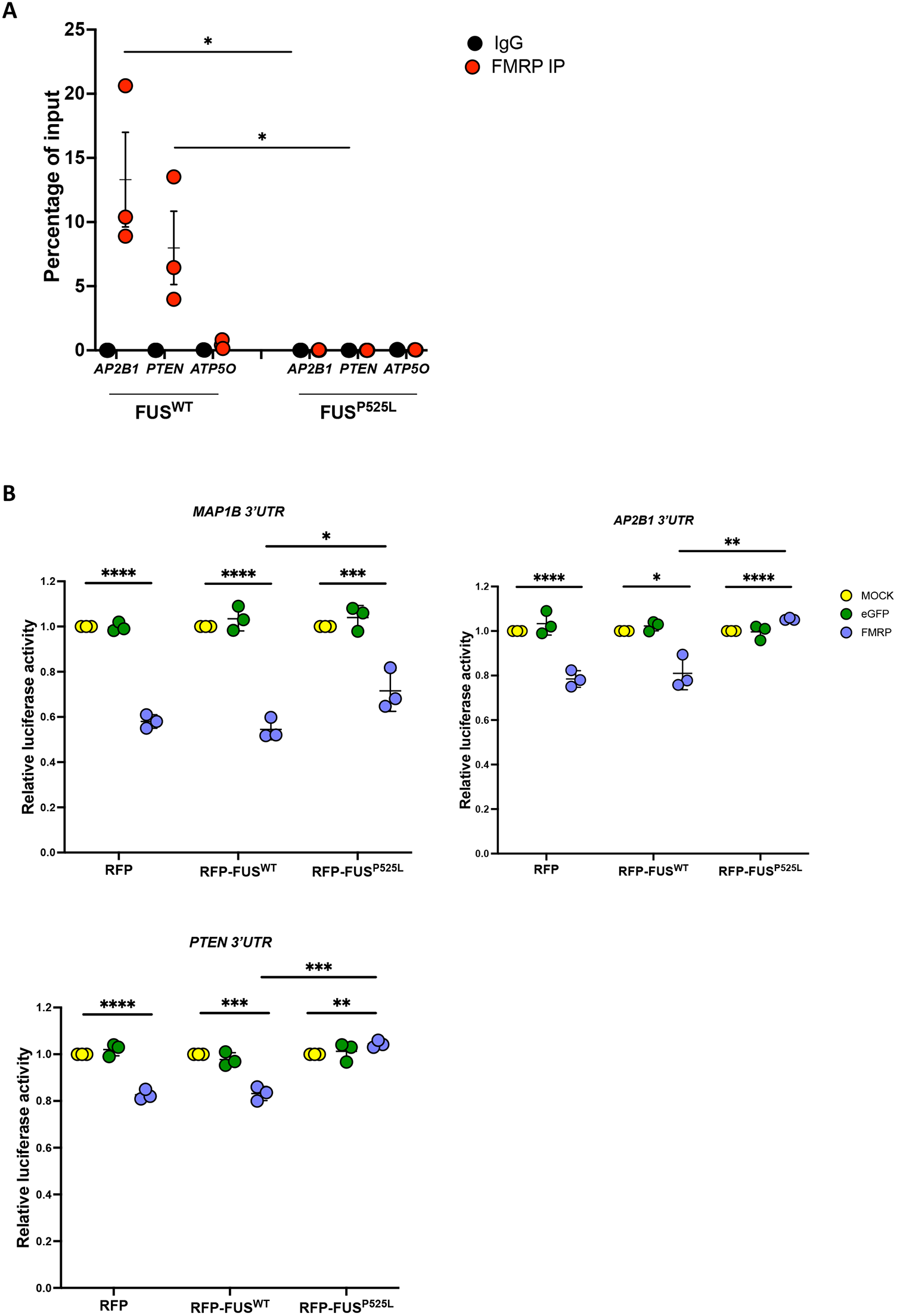
FMRP and FUS^P525L^ compete for *AP2B1, PTEN* and *MAP1B* 3’UTR binding. (**A**) Analysis of *AP2B1* and *PTEN* mRNA levels by real-time qRT-PCR in FMRP RIP samples from FUS^WT^ or FUS^P525L^ iPSC-derived spinal MNs. The housekeeping gene *ATP5O* is used as negative control. The graph shows the relative enrichment of the mRNAs pulled down by FMRP immunoprecipitation (IP), calculated as the percentage of input, in IP or control IgG samples, after normalization with an artificial spike RNA. The average from 3 independent differentiation experiments is showed and error bars indicate the standard deviation (Student’s t-test; unpaired; two tails; *p < 0.05). (**B**) Luciferase assay in HeLa cells expressing RFP, RFP-FUS^WT^ or RFP-FUS^P525L^ and transfected with Renilla luciferase reporter constructs containing the 3’UTR of *AP2B1* (RLuc-*AP2B1*-3’UTR), *MAP1B* (RLuc-*MAP1B*-3’UTR) or *PTEN* (RLuc-*PTEN*-3’UTR), in combination with plasmids overexpressing FMRP, eGFP or alone (MOCK) (Student’s t-test; paired; two tails; *p < 0.05; **p < 0.01; ***p < 0.001; ****p < 0.0001).

**Figure 7.**
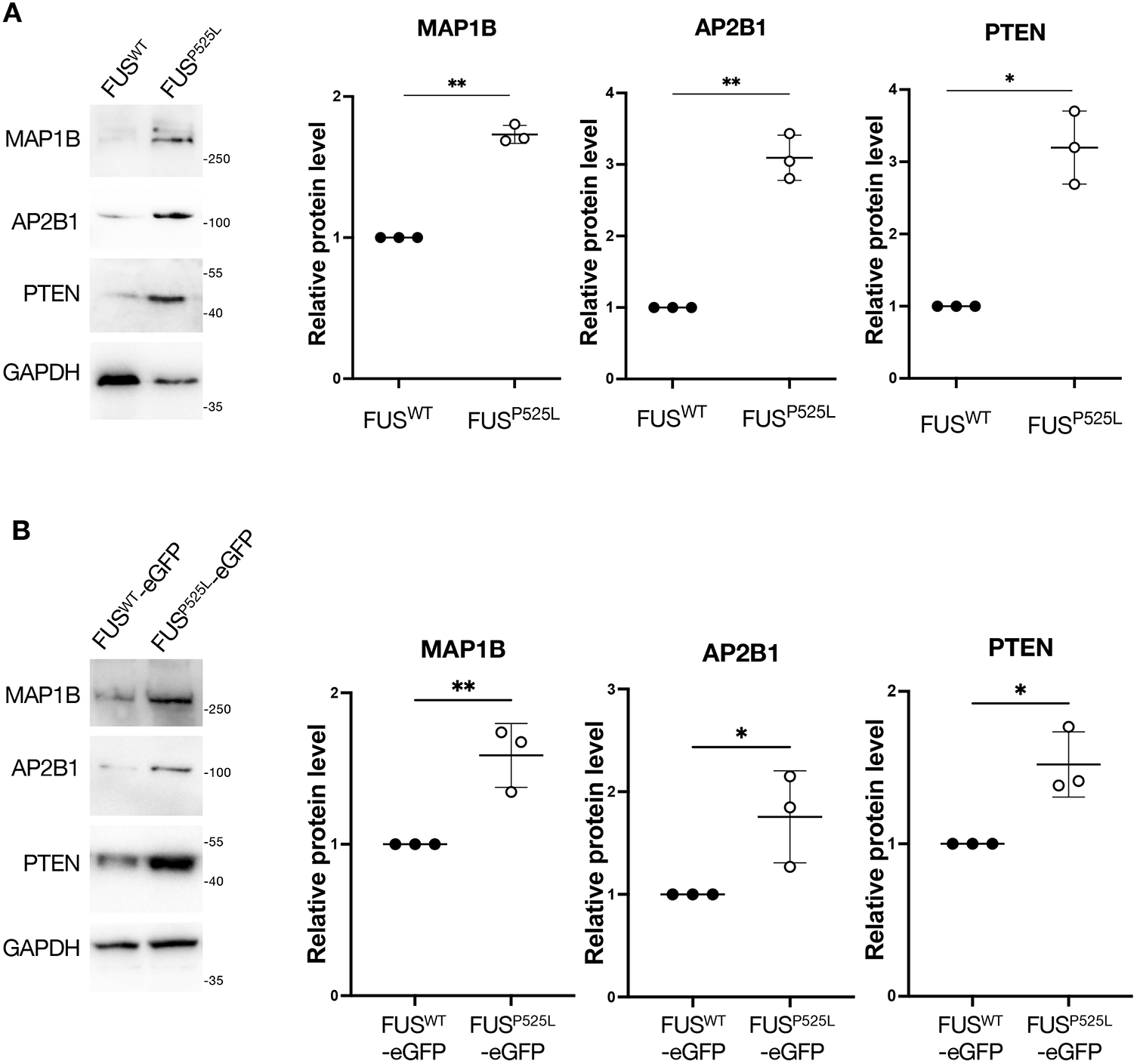
AP2B1, PTEN and MAP1B protein levels are increased in FUS mutant MNs. (**A**) Western blot analysis of the indicated genes protein levels in FUS^WT^ and FUS^P525L^ iPSC-derived spinal MNs. The molecular weight (kDa) is indicated on the right. The graphs show the average from three independent differentiation experiments, error bars indicate the standard deviation (Student’s t-test; unpaired; two tails; *p < 0.05; **p < 0.01). GAPDH signal was used for normalization. Protein levels are relative to the FUS^WT^ sample for each experiment. (**B**) Same analysis as in (A), performed in MNs from an independent isogenic pair of wild-type and P525L iPSC lines, in which endogenous FUS is fused at the C-terminal to the enhanced green fluorescent protein (eGFP) (Marrone et al., 2018).

Collectively, these results show that *AP2B1, MAP1B* and *PTEN* represent novel common FMRP and FUS targets whose levels are kept low by FMRP in normal conditions, and aberrantly increased in FUS mutant MNs.

## DISCUSSION

Neurons are specialized cells with a high level of polarization defined by the presence of subcellular compartments such as the cell body, dendrites, axons, and synapses. Each neuronal subtype is endowed with a specific and highly regulated set of transcripts and proteins in each of these cellular compartments, establishing the complex network of molecular pathways essential for neuronal functions. This architecture is fundamental for proper circuit formation and neuronal communication. Indeed, regulating mRNAs localization and local synthesis processes in separate subcellular compartments ensures an appropriate subcellular proteome, allowing neurons to respond to external cues and metabolic needs. Alteration of these mechanisms due to mutations, mislocalization and/or aggregation of neural RBPs leads to nervous system diseases. Identification of RBP targets and characterization of the mechanism by which this normal physiological state is perturbed in different subcellular compartments, in the cell type specifically affected by a given neurodegenerative disease, is of crucial importance. Moreover, to ensure proper RBP levels and activities, neurons rely on extensive auto- and cross-regulatory mechanisms. Thus, alteration on a node of such RBP network might lead to broader consequences on the neuronal RNA metabolism, as observed in complex diseases including neurodegeneration, epilepsy, and autism spectrum disorders. To this regard, FMRP and FUS, genetically linked to Fragile-X syndrome and ALS respectively, are two well-characterized multifunctional RBPs involved in several aspects of post-transcriptional gene expression. Notably, changes in the activity of FMRP in FUS-ALS models have been recently reported by us and others: FMRP might be caught into mutant FUS insoluble aggregates (Blokhuis et al., 2016); mutant FUS might increase FMRP phase separation by its recruitment in cytoplasmic ribonucleoprotein complexes (RNPs) (Birsa et al., 2021); and FMRP binding to 3’UTRs might be impaired by mutant FUS competition (Garone et al., 2021; this work). Collectively, these mechanisms could all contribute to compromise the proper functions of FMRP, inhibiting its role as a negative translation regulator with devastating effects on the RNA metabolism in MNs. Moreover, the incorrect cytoplasmic localization of mutant FUS could not only directly enrich the expression of some common targets through competition with FMRP, but indirectly promote non-physiological activities in other cellular compartments, an event often underlying brain disease.

We recently reported that one of the consequences of mutant FUS interference in FMRP functions is HuD upregulation (Garone et al., 2021). Here we show that HuD overexpression, in otherwise normal MNs, produces changes strikingly similar to those observed in mutant FUS MNs on the expression of a set of genes that are: a) involved in nervous system diseases; b) targets of wild-type FUS in intronic regions; c) targets of mutant FUS in the 3’UTR; d) targets of FMRP; e) involved in synaptic transmission and neuron development. This finding, together with previous evidence of HuD as a component of pathological cytoplasmic inclusions in FUS, TDP-43 and sporadic ALS patients (Blokhuis et al., 2016; De Santis et al., 2019), support the importance of this factor in the context of a complex regulatory RBP network, which is in place in normal MNs and disrupted upon FUS mutation. Notably, HuD upregulation was recently reported in the motor cortex of sporadic ALS patients devoid of mutations in known ALS genes (Dell’Orco et al., 2021). It will be interesting, in future studies, to assess the impact in MNs of genes commonly dysregulated by HuD overexpression and FUS mutation, identified in the present work.

In this work we found that three additional common FUS and FMRP targets might be dysregulated in ALS by the same competition mechanism proposed for HuD. Like HuD, regulation of *MAP1B, AP2B1* and *PTEN* translation by the two RBPs depends on their 3’UTR. Increased synaptic levels of MAP1B were previously reported in a FUS zebrafish model (Blokhuis et al., 2016). More recently, FUS-dependent upregulation of *MAP1B* has been found in cells carrying ALS-associated mutations in the Ubiquilin-2 (UBQLN2) gene (Strohm et al., 2022). Competition between FUS and FMRP for *MAP1B* mRNA binding at a G quadruplex structure in the 5’UTR has been previously proposed (Imperatore et al., 2020). Notably, reanalysis of our previous PAR-CLIP data (De Santis et al., 2019) suggested binding of mutant FUS to the 3’UTR of *MAP1B* and we show here that a reporter fused to the *MAP1B* 3’UTR is downregulated upon FMRP overexpression. This effect was reversed upon mutant FUS expression. Together, these data suggest that the competition between FUS and FMRP might occur at both UTRs of common target genes.

*AP2B1* encodes for a subunit of the AP2 adaptor, a complex involved in clathrin-mediated endocytosis, and its knockdown reduces the number of dendrites in developing neurons (Koscielny et al., 2018). To our knowledge, AP2B1 has never been studied in the context of neurodegeneration. However, increased levels of AP2B1 protein have been recently observed in the cerebrospinal fluid of Alzheimer’s disease patients (Sjödin et al., 2019). Our finding of increased levels of AP2B1 in FUS mutant MNs suggests that this protein might be explored as a possible biomarker in ALS as well.

PTEN, which is encoded by the *Phosphatase and tensin homolog* gene, has been extensively studied in the context of cancer and increasing evidence links this factor to neurodevelopmental and neurodegenerative diseases. The PTEN protein is part of a signaling network that contains multiple autism spectrum disorder (ASD)-associated gene products and represents a potentially common etiological mechanism for ASD and related neurodevelopmental disorders (Rademacher et al., 2019; Cummings et al., 2022). PTEN translation is negatively regulated by FMRP and heterozygous loss of *Pten* rescued neuronal phenotypes in a *Fmr1* knockout mouse (Sathyanarayana et al., 2022). Moreover, it has been shown that PTEN knockdown by RNA interference is beneficial for MN survival in SOD1- and C9ORF72-ALS and spinal muscular atrophy models (Kirby et al., 2011; Little et al., 2015; Stopford et al., 2017). Importantly, pharmacological inhibition of PTEN reduced MN loss and rescued neuromuscular innervation in a G93A SOD1 ALS model (Wang et al., 2021). Our data suggest that PTEN levels could be increased due to a strong impairment of FMRP binding and activity by mutant FUS, possibly extending the applicability of this therapeutic approach to FUS-ALS.

In conclusion, we propose that the analysis of common FMRP and FUS targets, which are dysregulated in soma and neurites of human MNs, could identify relevant biomarkers and therapeutic targets for ALS.

## MATERIALS AND METHODS

### Human iPSC culture and motor neuron differentiation

Human iPSC lines used in this study were previously generated and characterized: FUS^WT^ and FUS^P525L^ (WT I and FUS-P525L/P525L lines) (Lenzi et al., 2015); FMR1 KO iPSC line (Brighi et al., 2021); HuD overexpressing iPSCs (SYN1::HuD) (Garone et al., 2021); KOLF WT 2 and P525L16 (LL FUS-eGFP) (a kind gift of J. Sterneckert) (Marrone et al., 2018). As indicated in the original studies, informed consent had been obtained from all patients involved prior to cell donation. Cells were maintained in Nutristem XF/FF (Biological Industries) in plates coated with hESC-qualified matrigel (Corning) and passaged every 4-5 days with 1 mg/ml dispase (Gibco).

To obtain spinal MNs, iPSCs were stably transfected with epB-NIL, an inducible expression vector containing the Ngn2, Isl1, and Lhx3 transgenes (De Santis et al., 2018; Garone et al., 2019), and differentiated by induction with 1 µg/ml doxycycline (Thermo Fisher Scientific) in DMEM/F12 (Sigma-Aldrich), supplemented with 1× Glutamax (Thermo Fisher Scientific), 1× NEAA (Thermo Fisher Scientific) and 0.5× Penicillin/Streptomycin (Sigma-Aldrich) for 2 days and Neurobasal/B27 medium (Neurobasal Medium, Thermo Fisher Scientific; supplemented with 1× B27, Thermo Fisher Scientific; 1× Glutamax, Thermo Fisher Scientific; 1× NEAA, Thermo Fisher Scientific; and 0.5× Penicillin/Streptomycin, Sigma Aldrich), containing 5 µM DAPT and 4 µM SU5402 (both from Sigma-Aldrich) for additional 3 days. At day 5, MN progenitors were dissociated with Accutase (Thermo Fisher Scientific) and plated on Matrigel (BD Biosciences)-coated treated dishes or in modified Boyden chambers (Millicell Cell Culture Inserts) 6-well hanging inserts 1.0 µm PET. For soma and neurite compartments isolation, iPSC-derived MNs were grown in modified Boyden chambers for additional 7 days. On day 12, each inner cell culture was washed by replacing the medium with PBS w/o Ca^2+^/Mg^2+^. After the wash, the inner cell culture was replaced with PBS w/o Ca^2+^/Mg^2+^ and scraped using a cell lifter to collect the soma compartment. The cell bodies were centrifuged for 10 min at room temperature at 13,000 rpm. The total soma RNA was extracted with the RNA extraction kit (VWR International PBI) following the manufacturer’s instructions. For the neurites, the membrane was peeled off from the plastic scaffold by a cutter and pulled into a 2.0 ml tube. The membrane was lysed with TRK lysis buffer of the Micro Elute Total RNA Kit (VWR International PBI) and left rotating on a wheel for 20 min at room temperature. RNA was extracted with the Micro Elute total RNA Kit following the manufacturer’s instructions.

### Immunofluorescence

Cells were fixed in 4% paraformaldehyde (PFA) in PBS for 10 min at room temperature. The modified Boyden chambers were washed with PBS, permeabilized and blocked for 15 min using a solution of 0.5% BSA, 10% HRS, 0.2% Triton X-100 in PBS (all from Sigma-Aldrich). Anti-TUJ1 (for TUBB3 detection; 1:1000; T2200; Sigma-Aldrich; RRID:AB_262133) was diluted in a solution of 0.5% BSA, 10% HRS in PBS and incubated with the cells for 1 h at room temperature. Cells were then washed in PBS and incubated for 1 h at room temperature with the fluorescent conjugated secondary antibody anti-rabbit Alexa Fluor 488 produced in donkey (1:200; IS20015-1; Immunological Sciences); DAPI (1:2000; Sigma-Aldrich) was diluted in 0.5% BSA, 10% HRS in PBS and used to stain nuclei. Cells were washed with PBS and imaged using an inverted Zeiss LSM 780 microscope.

### Soma and Axon compartments RNA profiling

The nCounter custom code set used in this study was designed and synthesized by NanoString Technologies (NanoString Technologies, Seattle, WA). A total of 80 genes, including controls, were comprised in the code set (**Supplementary Table S4**). Briefly, 50 ng of soma or neurite total RNA was hybridized to the capture and reporter probe sets at 65°C for 20 h and applied to the nCounter preparation station according to the manufacturer’s instructions. Data were collected using the nCounter Digital Analyzer (NanoString) via counting individual fluorescent barcodes and quantifying target mRNA molecules present in each sample. For each assay, a high-density scan (555 fields of view-FOV) was performed on the nCounter Digital Analyzer.

### Bioinformatics analysis

The NanoString nSolver 4.0 software (https://www.nanostring.com/products/analysis-software/nsolver) was used to subtract background and normalize the RNA count data using the geometric means of the positive controls and the housekeeping genes *ATP5O* and *THYN*. The background subtraction and data normalization were performed for each independent class comparison. Expression for each gene was averaged across the three independent replicate samples. GraphPad Prism 6.0 (GraphPad Software) was used to determine differentially expressed genes; a Wilcoxon signed-rank test (for two conditions) or one-way ANOVA (for three conditions) was performed on the sample normalized data to determine statistical significance. Unsupervised hierarchical clustering and Gene ontology analysis of differentially expressed genes was performed using Qlucore Omics Explorer software and GO profiler tool, respectively.

PAR-CLIP reads and transitions were derived from published data (De Santis et al., 2019); only ratio of T to C transitions > 1 (comparing FUS^P525L^ vs FUS^WT^) were considered for the comparison with the FMRP HITS-CLIP data (Darnell et al., 2011). Paired Student’s t-test was used to detect the alternative and exclusive binding of FUS-WT and FUS^P525L^ on intron and coding sequences, respectively. Intersection between FUS PAR-CLIP and FMRP HITS-CLIP datasets was performed considering only genes that were detected in both experiments.

### RNA immunoprecipitation (RIP)

Human iPSC-derived MNs at day 12 of differentiation were lysed with PLB Buffer (5 mM MgCl^2^, 10 mM HEPES (pH 7.0), 150 mM KCl, 5 mM EDTA (pH 8), 0.5% NP-40, 2 mM DTT, with 100 U/ml RNAase inhibitor and 1× protease inhibitor cocktail), incubated for 5 min on ice and centrifuged 10 min at 4°C at 14,000×g. Protein concentration of the supernatant was then measured by Bradford assay and a volume containing 1 mg of proteins was diluted in NT2 Buffer (50 mM Tris pH 7, 150 mM NaCl, 0.5 mM MgCl^2^, 0.05% NP-40, 1 mM DTT, 20 mM EDTA pH 8 with 100 U/ml RNAse inhibitor and 1× protease inhibitor cocktail). Protein G-coupled dynabeads (Invitrogen) were washed in NT2 Buffer, incubated with 10 µg of anti-FMRP (ab17722; Abcam; RRID:AB_2278530) or rabbit monoclonal anti-human IgG antibody (ab109489; Abcam; RRID:AB_10863040) and left rotating on a wheel for 1 h at room temperature in NT2 Buffer. Beads were then washed in NT2 Buffer and incubated with the diluted lysates in a final volume of 500 µl. Binding was carried out at 4 °C with the samples rotating on a wheel for 2 h. Beads were then washed three times and resuspended in ice-cold NT2. An artificial spike RNA, i.e., an in vitro transcribed RNA fragment derived from the pcDNA3.1 plasmid, was added to the RNA fraction, which was then lysed with 250 µl of TRIzol (Invitrogen) and extracted according to manufacturer’s instructions. RNA was analyzed by real-time qRT-PCR with iTaq Universal SYBR Green Supermix (Bio-Rad Laboratories).

### Real-time qRT-PCR

Total RNA was retrotranscribed with iScript Supermix (Bio-Rad Laboratories) and analyzed by real-time qRT-PCR with iTaq Universal SYBR Green Supermix (Bio-Rad Laboratories). *ATP5O* was used as the internal calibrator. Primers sequences are listed in **Supplementary Table S6**.

### Luciferase assay

The 3’UTR of the genes of interest was cloned in the pSI-Check2 vector (Promega) using the primers listed in **Supplementary Table S6**, downstream the hRluc coding sequence. The resulting pSI-Check2-3’UTR constructs were transfected alone or in combination with epB-Bsd-TT-FMR1 or epB-Bsd-TT-eGFP in 5×10^4^ pre-seeded HeLa cells expressing RFP, RFP-FUS^WT^ or RFP-FUS^P525L^ (as described in (Garone et al., 2021)) in a 24-well plate using Lipofectamine 2000 (Life Technologies), following manufacturer’s instructions. Cells were harvested 24 h of post-transfection and RLuc and FLuc activities were measured by Dual Glo luciferase assay (Promega), according to the manufacturer’s protocol.

### Western blot

Western blot analysis was carried out using anti-AP2B1 (1:2000; 15690-1-AP; Proteintech; RRID:AB_2056351), anti-MAP1B (1:1000; 21633-1-AP; Proteintech; RRID:AB_10793666), anti-PTEN (1:2000; 22034-1-AP; Proteintech; RRID:AB_2878977), anti-GAPDH (1:2000; MAB-10578; Immunological Sciences) primary antibodies and HRP Donkey Anti-Mouse IgG (H+L) (IS20404; Immunological Sciences) and HRP Donkey Anti-Rabbit IgG (H+L) (IS20405; Immunological Sciences) secondary antibodies, with NuPAGE 4-12% Bis-Tris gels (Life Technologies) in MOPS-SDS buffer (Thermo Fisher Scientific). Signal detection was performed with the Clarity Western ECL kit (Bio-Rad). Images acquisition was performed with a Chemidoc MP (Bio-Rad), using the ImageLab software (Bio-Rad) for protein levels quantification. Uncropped blots are shown in **Supplementary Figure S2**.

### Statistical analysis

Statistical analysis, graphs, and plots were generated using GraphPad Prism 6 (GraphPad Software). As indicated in each figure legend, Student’s t-test or ordinary one-way ANOVA was performed, and data set are shown in dot plots indicating mean ± standard deviation (st.dev.). Sample size and the definition of replicates for each experiment is also indicated in the figure legends. RNA or protein expression was averaged across the three independent replicate samples.

### Materials availability statement

All data generated or analyzed during this study are included in the manuscript and supporting files; Source Data files have been provided for all figures in the manuscript.

## Supporting information

Raw Data

## ACKNOWLEGMENTS

The authors wish to thank all the members of the Rosa lab for helpful discussion and the Imaging Facility at Center for Life Nano- and Neuro-Science, Fondazione Istituto Italiano di Tecnologia, for support and technical advice. We are grateful to Jared Sterneckert (Technische Universität Dresden, Germany) for sharing their hiPSC lines. This work was partially supported by Sapienza University, Fondazione Istituto Italiano di Tecnologia and a grant from Istituto Pasteur Italia-Fondazione Cenci Bolognetti to A.R.

## AUTHOR CONTRIBUTIONS

Conceptualization, M.G.G., D.S. and A.R.; Formal analysis, Investigation, Methodology, M.G.G., and D.S.; Project administration and Supervision, A.R.; Writing-original draft, A.R. and M.G.G. All authors read and approved the final manuscript.

## COMPETING INTERESTS

The authors declare no competing interests.

## FIGURE LEGENDS

**Supplementary Figure S1.**
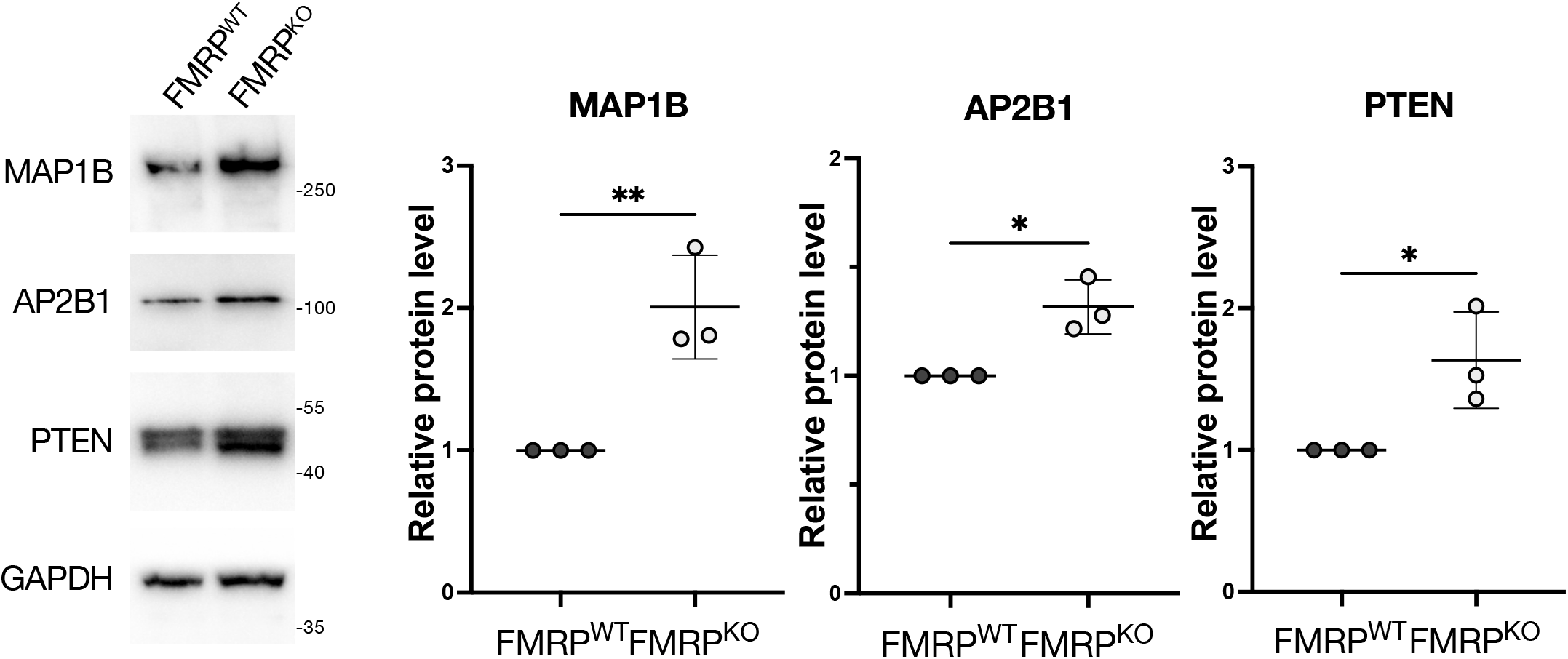
AP2B1, PTEN and MAP1B protein levels are increased in FMRP^KO^ MNs. Western blot analysis of the indicated genes protein levels in FMRP^WT^ and FMRP^KO^ iPSC-derived spinal MNs. The molecular weight (kDa) is indicated on the right. The graphs show the average from three independent differentiation experiments, error bars indicate the standard deviation (Student’s t-test; unpaired; two tails; *p < 0.05; **p < 0.01). GAPDH signal was used for normalization. Protein levels are relative to the FMRP^WT^ sample for each experiment.

**Supplementary Figure S2.**
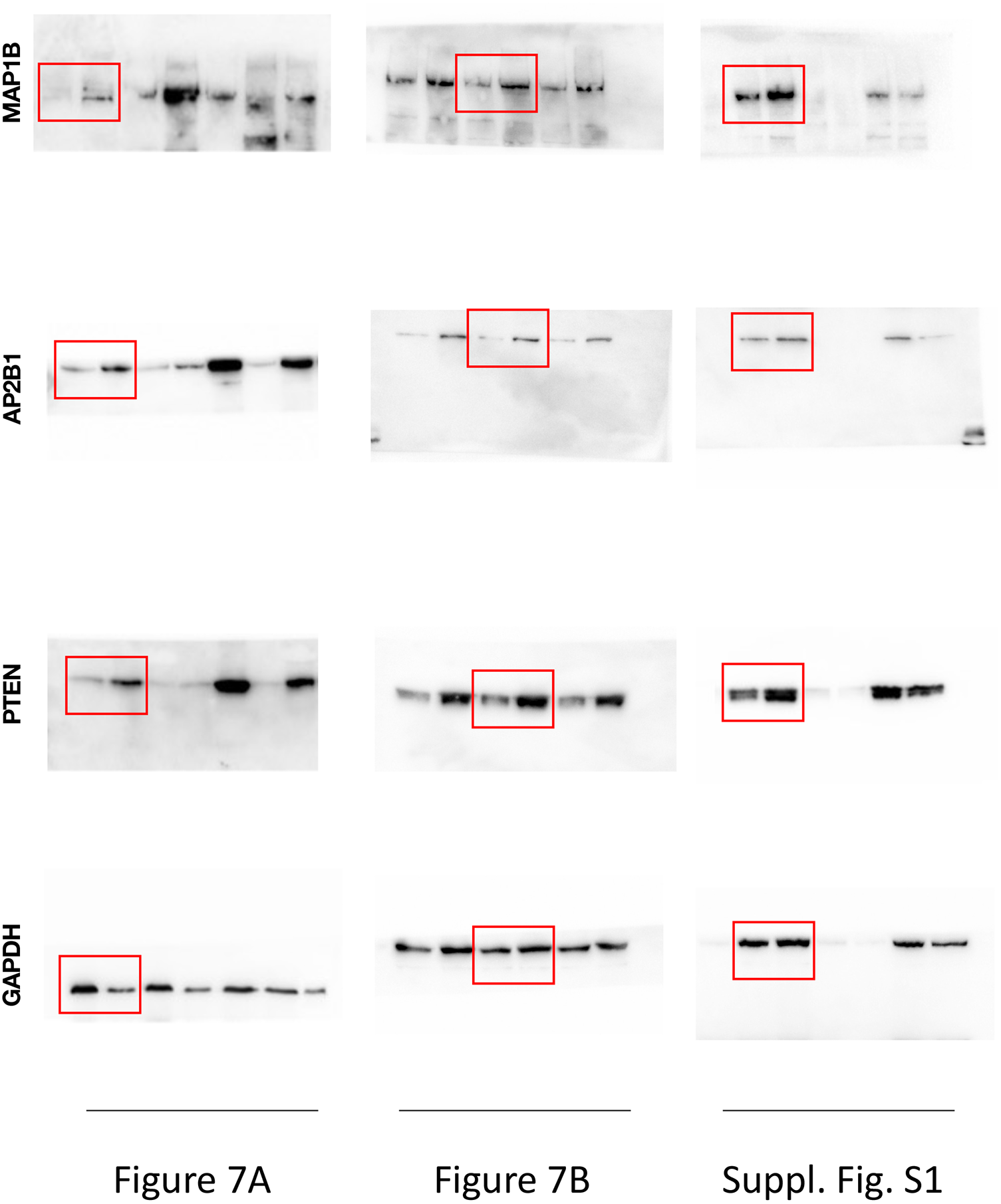
Uncropped western blot images. Uncropped western blot images relative to the experiments showed in Figure 7 and Supplementary Figure S1, indicated with a red box.

**Supplementary Table S1. Intersection between FUS and FMRP CLIP targets**

**Supplementary Table S2. GO term enrichment analysis of common FUS and FMRP targets**

**Supplementary Table S3. List of genes analyzed by digital color-coded molecular barcoding**

**Supplementary Table S4. nCounter custom code set used in this study**

**Supplementary Table S5. GO term enrichment analysis of differentially expressed genes**

**Supplementary Table S6. Sequences of the PCR primers**

## Notes

### Competing Interest Statement

The authors have declared no competing interest.

